# In vivo genome-wide CRISPR screens identify SOCS1 as a major intrinsic checkpoint of CD4^+^ Th1 cell response

**DOI:** 10.1101/2021.04.12.439455

**Authors:** Aurélien Sutra Del Galy, Silvia Menegatti, Jaime Fuentealba, Laetitia Perrin, Francesca Lucibello, Julie Helft, Aurélie Darbois, Michael Saitakis, Jimena Tosello, Derek Rookhuizen, Marc Deloger, Pierre Gestraud, Gérard Socié, Sebastian Amigorena, Olivier Lantz, Laurie Menger

## Abstract

The expansion of antigen experienced CD4^+^ T cells is limited by intrinsic factors. Using *in vivo* genome-wide CRISPR-Cas9 screens, we identified SOCS1 as a non-redundant checkpoint imposing a brake on CD4^+^ T-cell proliferation upon rechallenge. We show here that SOCS1 is a critical node integrating both IL-2 and IFN-γ signals and blocking multiple signaling pathways to abrogate CD4^+^ Th1 cell response. In CD8^+^ T-cell, SOCS1 does not impact the proliferation but rather reduces survival and effector functions. By targeting SOCS1, both murine and human CD4^+^ T-cell antitumor adoptive therapies exhibit a restored intra-tumor accumulation, proliferation/survival, persistence and polyfunctionality, promoting long term rejection of established tumors. These findings identify SOCS1 as a major intracellular checkpoint inhibitor of primed CD4^+^ T cells, opening new possibilities to optimize CAR-T cell therapies composition and efficacy.

## Introduction

Adoptive T cell therapy (ATCT) including T cells engineered with recombinant T Cell Receptor (TCR), Chimeric Antigen Receptor (CAR) or tumor-infiltrating lymphocytes (TILs) has become a powerful anti-cancer therapy. The *in vitro* manufacturing process enables to genetically reprogram a heterogenous mixture of CD4^+^ and CD8^+^ T-cell live drug to improve proliferation, survival and effector functions (Lim and June, 2017). Although CD8^+^ or CD4^+^ T cells alone can exert significant therapeutic effects (Adusumilli et al., 2014; Brentjens et al., 2003), the co-injection of both subsets is crucial for optimal and sustained antitumor activity (Borst et al., 2018; Linnemann et al., 2011; Sadelain, 2015). Exhibiting pleiotropic effects, CD4^+^ T cells can boost antitumor immune responses through both helper (Bos and Sherman, 2010; Corthay et al., 2005; Zhu et al., 2015) and cytotoxic functions (Kitano et al., 2013; Quezada et al., 2010; Śledzińska et al., 2020; Xie et al., 2010). However, after *in vitro* activation and adoptive transfer CD4^+^ and CD8^+^ T cells differ in their capacity to proliferate and persist *in vivo* (Turtle et al., 2016; Yang et al., 2017a). Hence, while CD8^+^ T cells undergo extensive and autonomous clonal expansion, CD4^+^ T cells need repeated antigen stimulation and rapidly stop to proliferate, leading to approximately 10-20 fold less expansion (Foulds et al., 2002; Homann et al., 2001; Ravkov and Williams, 2009; Seder and Ahmed, 2003). The differences in the magnitude and duration of their expansion are not due to external signals nor competition for resources (Homann et al., 2001; Seder and Ahmed, 2003). Instead, several studies reported that antigen experienced (Ag-exp) CD4^+^ T cells, including activated, effector and memory CD4^+^ T cells specifically curtail their own proliferation (Foulds et al., 2002; Helft et al., 2008; MacLeod et al., 2010; Merica et al., 2000). In the context of ATCT, as small doses of T cells are infused into patients, Ag-exp CD4^+^ T cells can become a limiting subset compromising an efficient protective immune response (Homann et al., 2001).

Using TCR-Transgenic (Tg) CD4^+^ T cells, we previously developed an *in vivo* system modeling a localized and asynchronous immune response, where new and returning T cells continuously enter the draining lymph node (Helft et al., 2008). We evidenced a preferential exclusion of Ag-experienced CD4^+^ T cells from an ongoing immune response. This inhibition is Ag specific, begins at day 2 (long before Ag disappearance) and is neither due to extrinsic factors, such as regulatory T cells (Tregs), lack of antigen presenting cell (APCs) education nor competition for Ag (Helft et al., 2008). Instead, Ag-experienced CD4^+^ T cells are stopped by an active and dominant phenomenon, which cannot be overcome by providing new Ag-loaded DCs. In this model, generalizable to several TCR-Tg CD4^+^ T cells, the expansion of Ag-exp CD4^+^ T cells is abolished while naive CD4^+^ T cells proliferation is maintained during the immune response. This strong, reproducible and intrinsic inhibition of Ag-exp CD4^+^ T cell proliferation allows for an *in vivo* efficient selective pressure of proliferative T cells after genetic modifications.

Using an *in vivo* genome-wide CRISPR-Cas9 positive screen, we interrogated in a systematic and unbiased manner the genes that restore the abrogated proliferation of Ag-exp Cas9 CD4^+^ T cells. Our screen identifies Suppressor of Cytokine Signaling 1 (SOCS1) as a non-redundant and intrinsic inhibitor of CD4^+^ T-cell proliferation and survival. In addition, we demonstrate that SOCS1 is a critical node, integrating cytokines signals (IFN-γ and IL-2) to actively limit CD4^+^ T cell functions. We investigated the function of SOCS1 in both mouse and human CD4^+^ and CD8^+^ antitumor adoptive cell therapies. SOCS1 inactivation restored CD4^+^ T-cell expansion, as well as helper and cytotoxic functions whereas it greatly boosted CD8^+^ T cell cytotoxic potential.

## Results

### In vivo genome-wide screen identified SOCS1 as a major non-redundant inhibitor of antigen experienced CD4^+^ T-cell expansion

To unravel the inhibitory mechanisms controlling the proliferation of Ag-exp CD4^+^ T-cell, we used the A^b^:Dby–specific Marilyn monoclonal CD4^+^ T cells (from the TCR-Tg *Rag2*^−/−^ Marilyn mouse (Lantz et al., 2000)). After intravenous (i.v.) adoptive transfer of naive CD45.2 Marilyn CD4^+^ T cells into C57BL/6 hosts, we initiated an immune response by injecting Dby peptide-loaded dendritic cells (DCs) into the footpad (**Fig. 1A**). To track the fate of newly recruited Ag-specific CD4^+^ T cells into such an ongoing immune response, we let the first cohort of primed Marilyn cells expand for a week. Then, by injecting i.v. a second cohort of naive or Ag-exp CD45.1 Marilyn CD4^+^ T cells (both effector and memory cells, generated *in vivo*), we previously demonstrated that the proliferation of Ag-exp CD4^+^ Marilyn T cells is strongly inhibited during an ongoing immune response while naive T cells are able to expand efficiently (Helft et al., 2008).

**Fig. 1.**
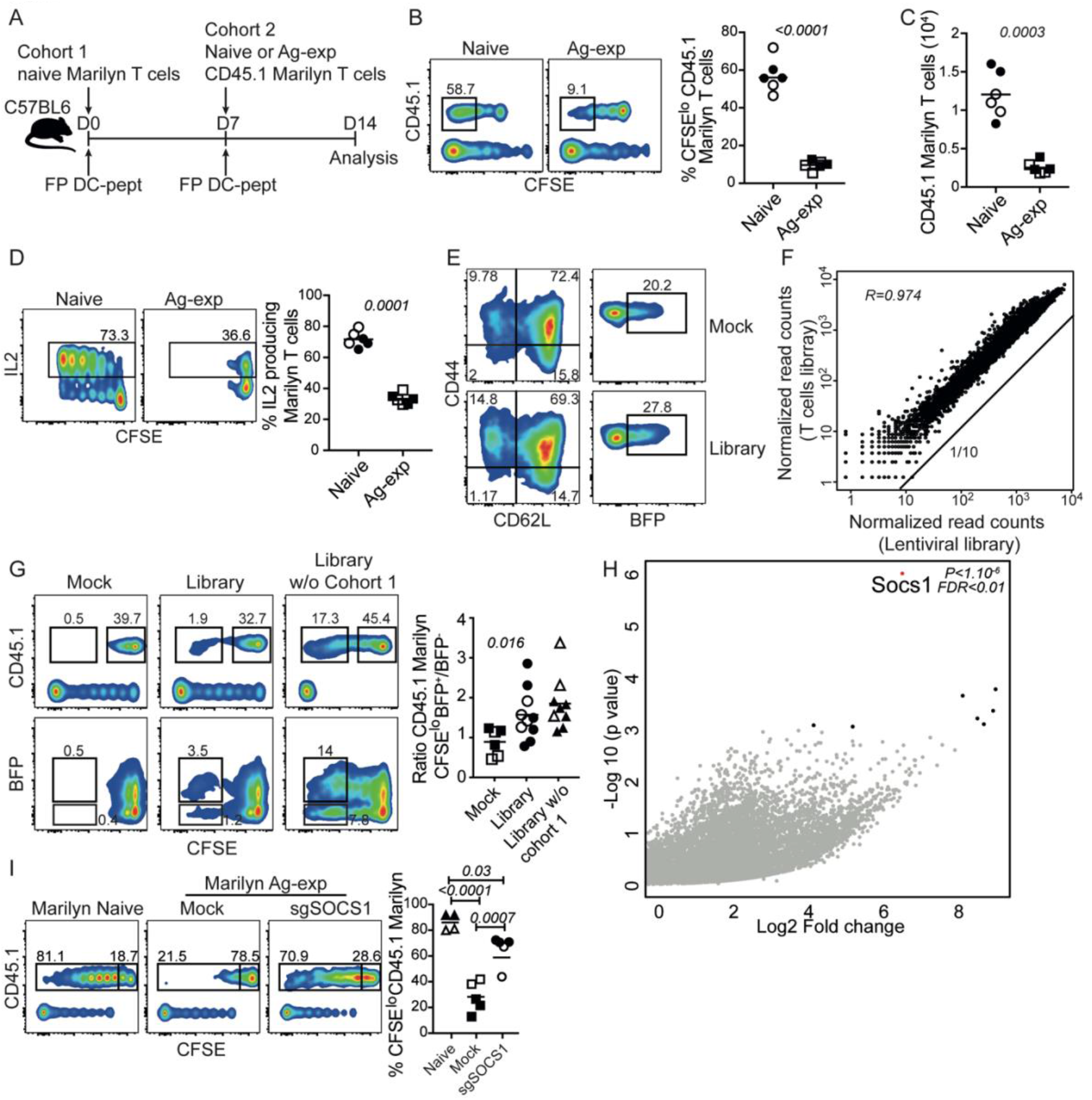
*In vivo* genome-scale (18400 genes) CRISPR pooled screens identify SOCS1 as non-redundant inhibitor of Antigen-experienced (Ag-exp) CD4 T cell expansion during an ongoing immune response. **(A)** Two cohorts experimental design to assess naive and Ag-exp CD4 T cell expansion in the course of an ongoing immune response used in **B-D**. (**B**) Flow plots and percentage (percentage highlighted are from singlets live CD45.1^+^ CD4 T cells) of proliferating Marilyn CD4 T cells, either 10^6^ naive or 2.10^6^ Ag-exp *in vitro* (based on CD62L positivity, reflecting a similar capacity to home to the LN), during an ongoing immune response in C57BL/6 mice. Mice were injected with 10^6^ cells intravenously and primed *in vivo* by injection of 10^6^ Dby peptide–loaded LPS-matured DCs into the footpad. (**C, D)** Survival and IL2 production of CD45.1 Ag-exp CD4 T cells compared to naïve CD45.1 Marilyn CD4 T cells during a recall response *in vivo*. (**E**) Ag-exp Cas9-Marilyn CD4 T cells CD44/CD62L phenotype and lentiviral library transduction efficiency (BFP^+^), prior to puromycin selection and injection *in vivo*. (**F)** Scatter plot comparing sgRNA normalized read counts in the original plasmid DNA library and in the transduced T cells after 4 days of puromycin selection (5μg/mL). (**G**) Representative flow plots and quantification of proliferating CD45.1-library-transduced Cas9-Marilyn CD4 T cells compared to CD45.1-Mock-transduced Cas9-Marilyn CD4 T cells, in the presence of the first cohort or not. Mice were injected with 12.10^6^ CD4 T cells IV and primed with 4.10^6^ Dby peptide–loaded LPS-matured DCs in the footpad at day 0 and day 7. (**H**) Enriched hits in the CFSE^lo^ subset of CD45.1-library-transduced CD4 T cells compared to the CFSE^hi^ subset in an ongoing immune response *in vivo* (strong selection). **(I)** Representative plots and percentage (gated on singlets live CD45.1^+^ CD4 T cells) of proliferating Ag-exp Mock Marilyn or sgSOCS1 Marilyn cells during a recall response, at day 14. Mice were injected with 10^6^ CD4 T cells IV and primed with 10^6^ peptide–pulsed LPS-matured DCs at day 0 and day 7. (G, H) Data shown are from two independents primary GW screens. (H) *p-value* corresponds to the gene-level enriched *p-value* and log2 fold change (LFC) to the median LFC of all sgRNA supporting the enriched RRA score. Targets with an FDR < 0.5 are highlighted in black. Each point is an individual mouse, open symbols are replicates from independent experiments (FP: footpad, DC: dendritic cells, pept: peptide, Ag-exp: antigen-experienced).

In this monoclonal recall response, we first reproduced the robust functional inhibition of Ag-exp CD45.1 Marilyn CD4^+^ T cells, generated *in vitro* by priming lymph nodes and splenocytes with Dby peptide, IL-2 and IL-7 and resting for 6-10 days (**Fig. 1B-D, Fig. S1A, B**). This model opens up the possibility to genetically manipulate Ag-exp CD4^+^ T cells before analyzing their fate *in vivo* during an immune response. To identify the intrinsic negative regulators of the CD4^+^ T cell immune response, we performed a positive genome-wide CRISPR screen looking for genes whose inactivation would restore the proliferation of Ag-exp CD4^+^ T cells during an immune response. We transduced *in vitr*o-generated Ag-exp Marilyn-R26-Cas9 (Cas9) T cells (**Fig. S1B**) with a genome-wide knockout (GWKO) sgRNA lentiviral library (18400 genes, 90K sgRNA)(Tzelepis et al., 2016), achieving 20-25% efficiency (BFP^+^) (**Fig. 1E**). After puromycin selection, 40% of the transduced T cells survived, revealing a 75% single infection rate (Chen et al., 2015) (**Fig. S1C**). Prior injection into the adoptive hosts, resting mock and library-transduced Ag-exp Marilyn-Cas9 T cells exhibited a central memory phenotype (CD62L^+^CD44^+^), allowing them to similarly home to the dLN (**Fig.1E**). Analysis of sgRNA in the transduced Marilyn-Cas9 T cells revealed that less than 0.5% of the sgRNA were under-represented as compared to the original plasmid library (**Fig. 1F, Fig. S1D, E**). Consistent with the coverage of sgRNAs *in vivo*, we conducted two independent GWKO pooled screens (*n=10 mice*) by injecting 12.10^6^ Ag-exp library-transduced or 12.10^6^ mock-transduced Marilyn-Cas9 T cells per C57BL/6 mouse as a second cohort (>130 Marilyn cells/gRNA/mouse with 5 different gRNA/gene e.g We next assessed the impact of SOCS1 >600 Marilyn cells/mouse with a mutated gene) (**Fig. 1A**). Seven days after transfer and priming, mock-transduced Marilyn-Cas9 cells proliferation was abolished. However, the proliferation of the library-transduced Marilyn-Cas9 cells in the presence of the first cohort (strong *in vivo* selection) was significantly restored, as shown by the higher ratio of BFP^+^/BFP^-^ in the CFSE^lo^ subset compared to mock-transduced Marilyn-Cas9 cells ratio, indicating the release of the proliferative blockade by some sgRNA (**Fig. 1G**). Without the first cohort, library-transduced Marilyn cells expanded to some extent, attesting for an efficient priming (**Fig. 1G**). After CFSE-based cell sorting of CD45.1 Marilyn-Cas9 T cells (**Fig. S1F**), amplified sgRNA sequences enriched in the CFSE^lo^ subset were compared to sgRNA from non-dividing T cells (CFSE^hi^). The small fraction of sgRNA represented in the CFSE^lo^ subset attest the effectiveness of the *in vivo* selection (**Fig. S1G**). Analysis of individual sgRNA enriched in the CFSE^lo^ subset of two independents screens from the strong *in vivo* selection identified *Socs1* as the major gene involved in the restored proliferation of Ag-exp CD4^+^ T cells *in vivo* (*p*<1.10^−6^, false discovery rate (FDR) <1%) (**Fig. 1H**), while other lower ranking targets presented a FDR >0.5. Interestingly, *Socs1* sgRNA were also significantly enriched in the CFSE^lo^ subset of library-transduced Marilyn cells injected compared to the CFSE^hi^ subset (weak *in vivo* selection), consistent with the capacity of Ag-exp CD4^+^ T cells to inhibit one another (**Fig. S1H**). Altogether these data support a non-redundant and critical role for SOCS1 in T cell biology, in particular for CD4^+^ T cells that had not yet been explored.

We next assessed the impact of SOCS1 inactivation on Ag-exp CD4^+^ T-cell proliferation using electroporation of individual sgRNA Cas9 ribonucleoprotein complexes (RNPs) (Seki and Rutz, 2018) in two different CD4^+^ TCR-Tg models, Marilyn, and OT2 cells (the latter expresses a TCR specific for MHC-II restricted ovalbumin peptide). Briefly, *in vitro* primed CD4^+^ TCR-Tg cells were electroporated with RNPs, comprising a sgRNA targeting a different sequence in *Socs1* gene than those from the GWKO library. 10^6^ naïve, 2.10^6^ Ag-exp mock or 2.10^6^ Ag-exp sgSOCS1 CD4^+^ T cells (based on CD62L positivity) (**Fig. S1 B, I, J**) were CFSE-labeled and subsequently injected as secondary responders into C57BL/6 mice during an ongoing immune response. In both models, the large naïve CD4^+^ T cell expansion indicated efficient priming whereas *Socs1* gene inactivation unleashed the brake observed in mock Ag-exp CD4^+^ T cells proliferation (**Fig. 1I, Fig.S1K**). These results uncover a role for SOCS1 as a major intrinsic regulator responsible for Ag-exp CD4^+^ T cell arrest during an ongoing immune response. Notably, we did not observe any Treg conversion after Marilyn and OT2 cells transfer *in vivo* (**Fig. S1L**), suggesting that Ag-specific Tregs are not involved in our models, contrary to what was suggested in another report (Akkaya et al., 2019).

### SOCS1 is a critical node integrating multiple cytokine signals to actively inhibit CD4^+^ T cell functions

To mechanistically characterize SOCS1-mediated inhibition of CD4^+^ T cells, we sought for potential inducers and subsequently assessed the functional consequences of *Socs1* inactivation on Ag-exp CD4^+^ T cells. SOCS1 expression in murine splenocytes and CD4 T cells is induced by both cytokines and TCR stimulation with different timelines and intensities (Sukka-Ganesh and Larkin, 2016a). Although basal levels of SOCS1 are present in untreated T cells, increase in SOCS1 protein level in response to cytokine stimulation arises rapidly (6 hours) while its maximal expression occurs 48h after TCR stimulation (Egwuagu et al., 2002; Sukka-Ganesh and Larkin, 2016b). This is in accordance with the timeframe of inhibition in our model, which starts *in vivo* 2 days after priming (Helft et al., 2008). These results suggest that TCR engagement could be the reason for SOCS1 induction in Ag-exp CD4^+^ T cells. To assess if a differential sensitivity to cytokine signaling could explain the selective inhibitory activity between naïve and antigen experienced cells, we compared the transcriptional expression of cytokine receptors between sorted proliferating (green) and inhibited subsets (red) during an ongoing immune response, at day 14 (**Fig. 2A**). We observed a significantly increased expression of *Il2ra* (also called CD25, confirmed at protein level, **Fig. S2A***),Ifngr1* and *Ifngr2* in the CFSE^hi^ cells as compared to CFSE^lo^ cells (**Fig. 2B**). Moreover, naive and Ag-exp CD4^+^ T cells secreted IL-2, while only Ag-exp Marilyn CD4^+^ T cells produced both IL-2 and IFN-γ (**Fig. S2B**). As SOCS1 is a known regulator of IFN-γ signaling (Alexander et al., 1999), we evaluated the proliferation of Ag-exp IFN-γR^−/−^ Marilyn cells during an ongoing immune response, but the absence of the receptor marginally restored the expansion of these cells *in vivo* (**Fig. 2C**). SOCS1 can also be induced by IL-2 in T cells and associates with IL-2Rβ (Liau et al., 2018) to potently inhibit IL-2–induced Stat5 function (Sporri et al., 2001). Using blocking antibodies concomitant with Ag re-stimulation of Ag-exp Marilyn CD4^+^ T cells *in vivo*, we then assessed the roles of IL-2 and IFN-γ alone and in combination in this inhibition. The blockade of IL-2 signaling using anti-mouse IL-2Rβ, which inhibits binding of IL-2 to the IL-2R did not reverse Ag-exp CD4^+^ T cells impaired proliferation (**Fig. 2D**). However, blockade of both IL-2 and IFN-γ signaling (using anti-IFN-γRα and Ag-exp IFNγ-R^−/−^ Marilyn T, **Fig. S2C**) significantly rescued the expansion of re-stimulated Ag-exp Marilyn T cells (**Fig. 2D**). This shows a redundancy between the two cytokine receptors upstream of SOCS1 to impair Ag-exp CD4^+^ T cells expansion.

**Fig. 2.**
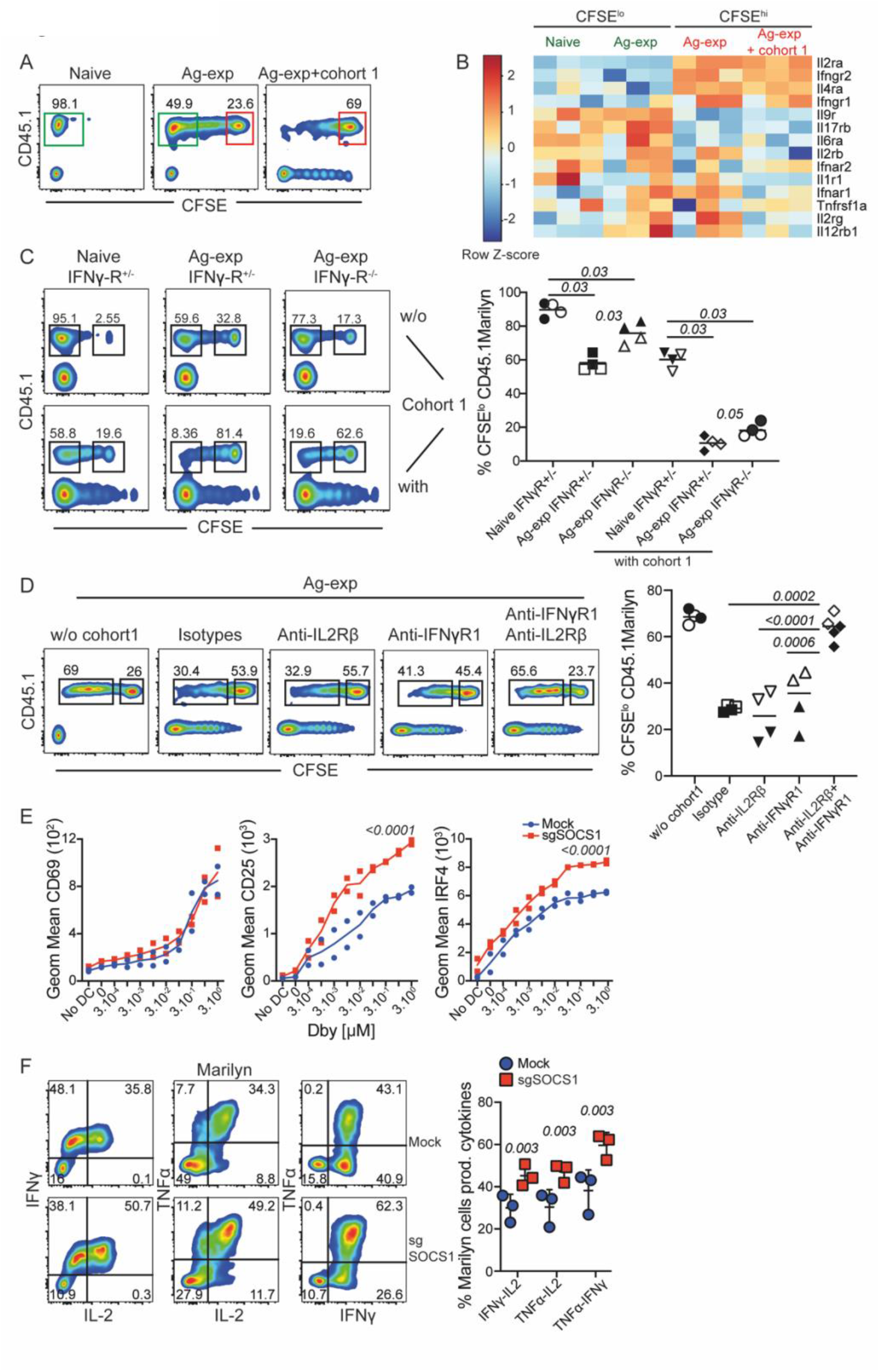
SOCS1 is a node integrating several cytokines signals to actively silence polycytokine release. (**A**) SORTing strategy of CFSE^lo^ (green) and CFSE^hi^ (red) naïve or Ag-exp Marilyn cells from an ongoing immune response. (**B**) Heat map displaying the expression of a selected list of cytokine receptors by proliferating or inhibited Marilyn cells (first seven receptors *p<0*.*01*, FDR<0.5). (**C**) Representative flow plots (percentage highlighted are from singlets live CD45.1^+^ CD4 T cells) and quantification of 10^6^ Marilyn naïve IFNγ-R^+/−^ or Marilyn Ag-exp IFNγ-R^+/−^ or Ag-exp IFNγ-R^−/−^ expansion *in vivo* after cells transfer and footpad vaccinations at day 14, with or without (w/o) cohort 1 expansion. (**D**) Representative flow plots (percentage highlighted are from singlets live CD45.1^+^ CD4 T cells) and quantification of 10^6^ Marilyn Ag-exp expansion *in vivo* during a recall response, in the presence of blocking antibodies (200μg) injected intraperitoneally at day 7, day 9, day 11: isotypes, anti-IL2Rβ, anti-IFNγRα. (**E**) Flow cytometric evaluation of CD69, CD25, IRF4 and expression in sgSOCS1 Ag-exp Marilyn compared to Mock cells after overnight co-culture with peptide–pulsed LPS-matured DCs *in vitro*, in the absence of cytokine. (**F**) Flow plots and percentage of IFN-γ-, TNFα- and IL-2-producing Mock or sgSOCS1 Marilyn. Values are shown as means or means ± SD. Each point is an individual mouse, open symbols are replicates from independent experiments, analyzed by Mann– Whitney U tests or two-way ANOVA (E).

Then, we estimated the functional consequence of *Socs1* deletion on Ag-exp CD4^+^ T cells TCR-induced activation, reflected by expression of the early activation marker CD69, the late activation marker CD25 and the T cell receptor responsive transcription factor Interferon Regulatory Factor 4 (IRF4) (**Fig. 2E, Fig. S2D, E**). After overnight stimulation with titrated peptide-pulsed DCs, both Ag-exp *Socs1*-inactivated Marilyn and OT2 cells displayed similar sensitivity (Ag dose leading to 50% of the maximum response) to Ag stimulation as compared to mock-treated cells. However, we observed a striking increase in CD25 and IRF4 expressions at higher Ag doses with an elevated “plateau” (**Fig. 2E, Fig. S2D, E**). This suggests that SOCS1 does not directly regulate proximal signals induced by cognate peptide stimulation but rather inhibits downstream signaling events. This would suggest the release of a negative feedback loop, related to the secretion of IL-2 and IFN-γ in the medium. As IRF4 is a the central regulator of Th1 cytokines secretion in CD4^+^ T cells (Mahnke et al., 2016; Wu et al., 2017), we evaluated the capacity of *Socs1* inactivated CD4^+^ T cells to display polyfunctionality. *Socs1* inactivated Marilyn and OT2 cells exhibited higher percentage of Th1 polycytokine (IFN-γ-, TNFα- and IL-2-) production after re-stimulation (**Fig. 2F, S2F**). Thus, by integrating several cytokine signals, SOCS1 actively hampers polyfunctionality of Ag-exp CD4^+^ T cells. Our findings show that SOCS1 is a node capable of receiving signals from several inputs (IFN-γ and IL2) to abrogate multiple signaling outputs, leading to blockade of proliferative and effector functions.

### Socs1-inactivation improves the intrinsic and extrinsic antitumor effect of adoptively transferred CD4^+^ T cells

The restored functionalities of *Socs1* inactivated CD4^+^ T cells led us to evaluate the direct and indirect therapeutic potential of *Socs1* deletion on adoptively transferred antitumor CD4^+^ T cells. We challenged female C57BL/6 mice with the Dby-expressing MB49 male bladder carcinoma cells and 10 days later intravenously transferred mock or sgSOCS1 antigen experienced Marilyn cells (**Fig. 3A**). In the absence of Marilyn cell transfer, the immunogenic but nevertheless aggressive MB49 tumors grew unimpeded by the endogenous immune response (**Fig. 3B**). The transfer of mock Ag-exp Marilyn cells led to rejection of MB49 tumors in 4 out of 11 mice, while the transfer of Ag-exp sgSOCS1 Marilyn CD4^+^ T cells induced tumor rejection in 9 out of 11 mice (**Fig. 3B, C**). To determine the mechanisms of this protection, we analyzed the number, phenotype and transcriptome of the transferred Marilyn T cells in the tumor, in the tumor draining lymph node (TdLN) and in a distant irrelevant LN (irr-LN). Seven days after transfer, the number of Ag-exp sgSOCS1 Marilyn cells was much higher in the tumor and TdLN as compared to mock Marilyn cells (**Fig. 3D**). This was associated with a higher percentage of proliferating Ag-exp sgSOCS1 Marilyn cells in TdLN-infiltrating as compared to mock Marilyn cells, which displayed dominant arrest in their proliferation (**Fig. 3E**). In addition to enhanced proliferation or survival of CD4^+^ T cells, bulk RNAseq analysis of Marilyn cells sorted from TdLN at day 7 showed that *Socs1* deletion increased the expression of cytokine receptors, *Il12rb2*, and *Il2rb*, effector molecule such as *Tbx21*, activation markers *Cxcr3* (Rabin et al., 2003), *Icos* and anti-apoptotic genes, such as *Hopx* (Albrecht et al., 2010) and *Pif1* (Gagou et al., 2011) (**Fig. S3A)**. Hallmarks analysis revealed that several pathways were significantly upregulated in sgSOCS1 TdLN infiltrating cells (*FDR < 0*.*05*). They include genes implicated in cell cycle and DNA replication (G2M checkpoints, E2F transcription factors, mitotic spindle) as well as IL2/STAT5 signaling (**Fig. 3F**). This mirrors the higher persistence of Ag-exp sgSOCS1 Marilyn cells as compared to mock Ag-exp Marylin in the blood of tumor challenged mice, 25 days after transfer (**Fig. S3B**). Analysis at the protein level of Ag-exp sgSOCS1 Marilyn T cells in the TdLN and in the tumor confirmed our bulk RNA-seq analysis with preserved expression of Th1 cytokines and cytotoxic molecules (Granzyme B) (**Fig. 3G, H, Fig. S3C**). However, the absence of infiltrating mock Marilyn T cells does not allow us to evalute if *Socs1* inactivation increase GzmB expression or rather increased the survival of sgSOCS1 Marilyn T cells with a preserved GzmB expression. Consistent with this functional analysis, major differences emerged in transcriptomic profiles related to T cell function and differentiation. Gene set enrichment analysis (GSEA) revealed downregulation of naive-associated genes and enrichment of T conventional marker (Tconv) as compared to Treg-related genes in Marilyn sgSOCS1 cells (**Fig. S3D**).

**Fig. 3.**
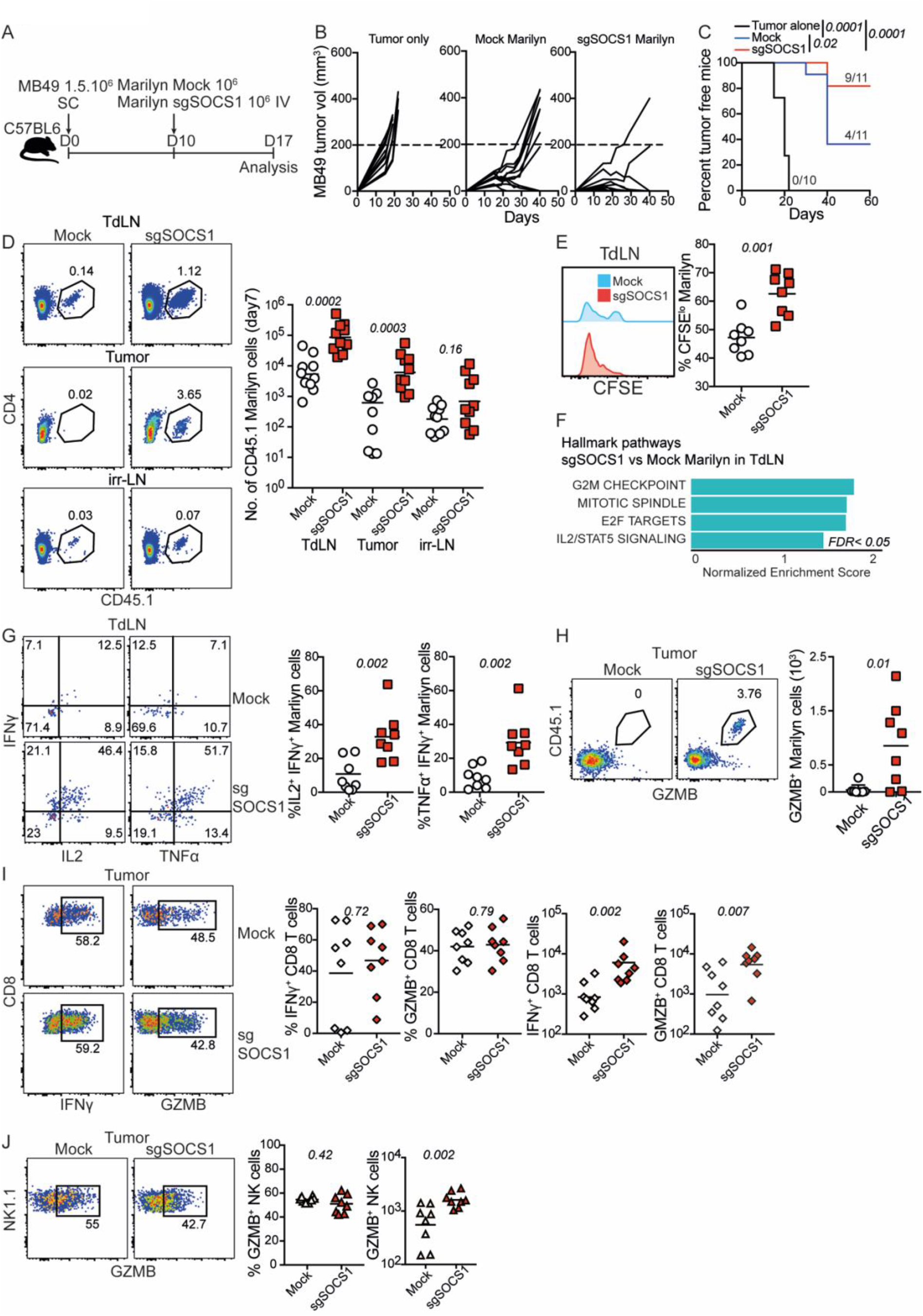
*Socs1* inactivation in Ag-exp Marilyn CD4 T cells enhances the rejection of male bladder MB49 tumors. (**A**) Schematic of Marilyn CD4 T cells (ACT) in C57BL/6 female mice-bearing the male DBY-expressing bladder tumor line MB49. (**B**) Growth curves of MB49 tumors in C57BL6 mice following the different ACT: PBS control, adoptive transfer of 10^6^ mock Ag-exp Marilyn or 10^6^ sgSOCS1 Ag-exp Marilyn Cas9. (**C**) Tumor-free survival following ACT, log-rank (Mantel–Cox) test. (**D**) Representative flow plots and quantification of Mock or sgSOCS1 Marilyn cells in the tumor draining lymph node (TdLN), in the tumor and in the irrelevant lymph nodes (irr-LN) at day 7 after ACT. (**E**) Representative flow plots and percentage of mock and sgSOCS1 Marilyn cells proliferation in the TdLN at day 7 after ACT. (**F**) Gene set enrichment analysis (GSEA) of selected hallmarks transcriptional signatures (MSigDB) with an FDR value < 0.05 in Ag-exp sgSOCS1 versus Ag-exp mock Marilyn T cells in the TdLN (n = 3 replicates from 2 pooled mice). (**G**) Representative flow plots and quantification of IFNγ^+^ IL2^+^ and IFNγ^+^ TNFα+-producing mock or sgSOCS1 Marilyn CD4 T cells in the TdLN at day 7 after transfer. (**H**) Flow plot and quantification of granzyme B (GZMB) expressed by tumor-infiltrating sgSOCS1 Marilyn CD4 T cells at day 7. (**I, J**) Representative flow plots and quantification of CD8- and NK-tumor infiltrating cells, producing effector molecules at day 7 after Marilyn cells transfer. Data are shown as mean, analyzed by Mann–Whitney U tests, from two independent experiments, *n=4-6* mice/group.

Importantly, the number of activated host polyclonal CD8^+^ T cells and NK cells in the tumor was increased by 2-3-fold in MB49-bearing mice transferred with sgSOCS1 CD4^+^ Marilyn cells, at day 7 (**Fig S3. E**). As estimated by *ex vivo* IFN-γ or GZMB expressions, the transfer of Ag-exp sgSOCS1 Marilyn T cells lead to increased number of functional effector cells at the tumor site (**Fig. 3I, J, Fig. S3E**). Thus, *Socs1* deletion in CD4^+^ T cells improved the anti-tumor response through both extrinsic and intrinsic mechanisms with enhanced CD4^+^ T cell expansion, function and persistence as well as increased magnitude of the endogenous anti-tumor immune response.

### Differential effect of Socs1-inactivation on the properties of CD4^+^ and CD8^+^ T cells used for adoptive transfer against melanoma tumors

To compare the biological impact of *Socs1* deletion in CD4^+^ and/or CD8^+^ T cells on anti-tumor response, we independently generated *in vitro* activated tumor specific CD4^+^ and CD8^+^ T cells in which we deleted or not SOCS1 as described above (**Fig. S4A, B**). We used CD90.1 OT2 CD4^+^ and CD45.1 OT1 CD8^+^ T cells recognizing MHC-II and MHC-I restricted ovalbumin peptides, respectively and subcutaneously implanted B16-OVA melanoma cells as tumor model, without conditioning or cytokines supply (**Fig. 4A**). As compared to the results displayed in Fig. 3, the inactivation of *Socs1* in OT2 cells had a marginal antitumor effect (**Fig. 4B, C**). This could be related either to the use of the highly immunosuppressive B16 melanoma model or to the co-transfer of large number of high avidity antitumor specific CD8^+^ T cells. However, after adoptive transfer of sgSOCS1 OT1 T cells (**Fig. 4B, C**), we observed a significant and durable rejection of established tumors as compared to transfer of mock OT1 T cells (*p<0*.*001*, log-rank). The infiltration of T cells seven days after transfer showed an increased accumulation in the TdLN and in the tumor for the group receiving both sgSOCS1 OT1 and sgSOCS1 OT2 cells as compared to mock transferred cells (**Fig. 4D**). Importantly, in the TdLN, *Socs1* inactivation had a profound effect on OT2 CD4^+^ T-cell proliferation with a large increase in fully divided CD4^+^ T cells, whereas the pattern of OT1 CD8^+^ T-cell proliferation was barely affected, suggesting that SOCS1 impacts CD8^+^ T-cell survival more than proliferation (**Fig. 4E**). However, as both OT1 mock and OT1 sgSOCS1 extensively proliferate, our design does not allow us to observe a significant difference in the CFSE profiles after 8 divivions.

**Fig. 4.**
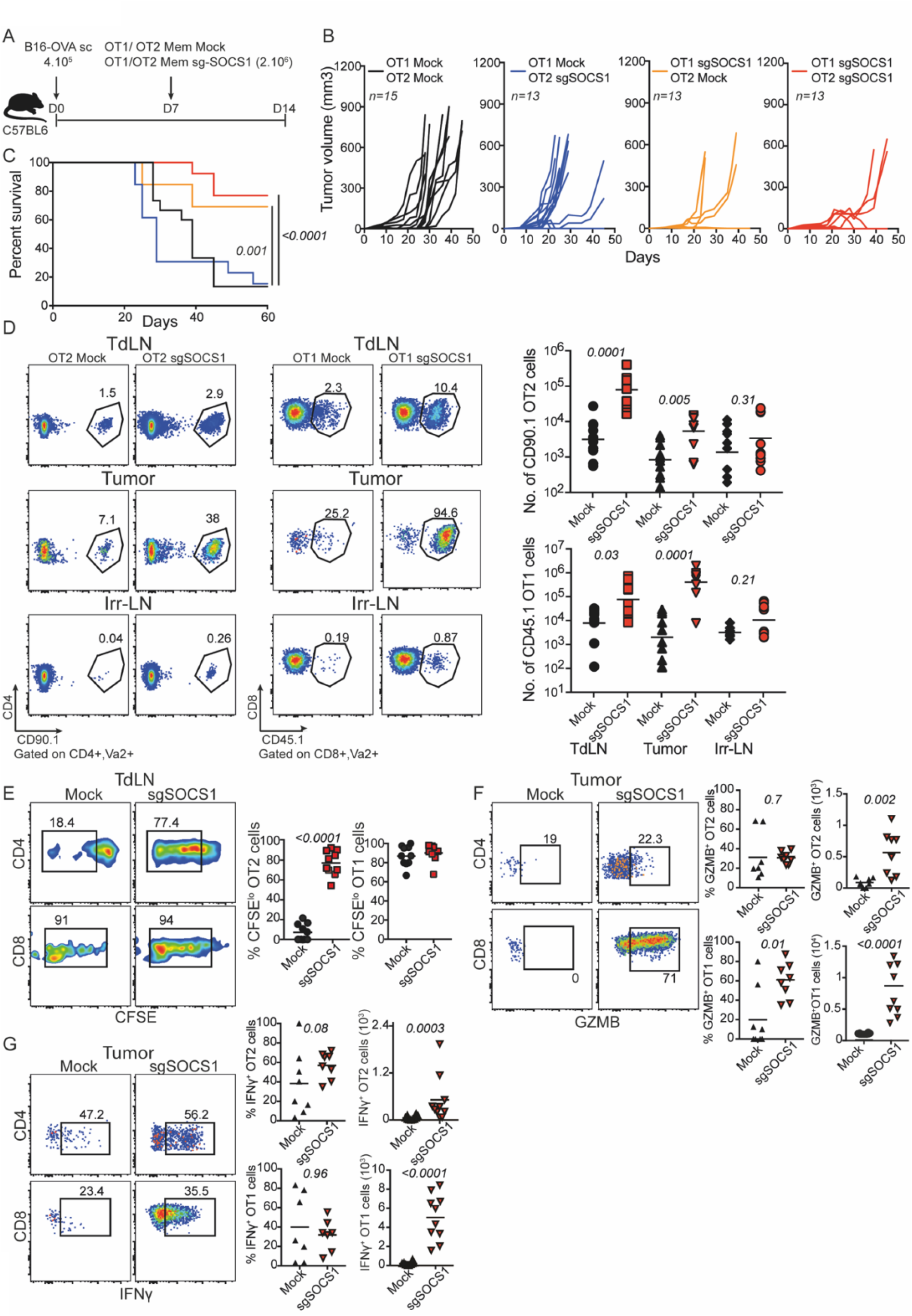
B16-OVA tumor rejection with improved ACT: *Socs1* gene inactivation restores the proliferation of OT2 cells and enhances OT1 cell survival and cytotoxicity. (**A**) Schematic of OT1 CD8- and OT2 CD4-adoptive T cell therapy (ACT) in C57BL/6 mice-bearing B16-OVA melanoma tumors. (**B**) Growth curves of B16-OVA tumors in C57BL6 mice following adoptive transfer with OT1 (2.10^6^ Mock or 2.10^6^ sgSOCS1) and OT2 cells (2.10^6^ Mock or 2.10^6^ sgSOCS1). (**C**) Kaplan-Meier survival analysis of B16-OVA-bearing mice following ACT, log-rank (Mantel–Cox) test. (**D**) Representative plots and quantification of Mock or sgSOCS1 OT1 and OT2 cells in the tumor draining lymph node (TdLN), in the tumor or in the irrelevant lymph nodes (Irr-LN) at day 7 after ACT, gated on singlets live Vα2+ T cells. (**E**) Representative flow plots and percentage of Mock or sgSOCS1 OT1 and OT2 cells proliferating in the TdLN at day 7. **(F, G**) Representative flow plots and quantification of mock or sgSOCS1 OT2 and OT1 tumor-infiltrating cells producing IFN-γ and granzyme B molecules and at day7 after transfer. Data are shown as mean, analyzed by Mann–Whitney U tests, from two independent experiments, *n=5-8* mice/group.

Sixty days after transfer, the number of sgSOCS1 OT2 cells ultimately decreased in the blood of B16-OVA challenged mice, while a population of central memory sgSOCS1 OT1 cells remained 15-fold more abundant than mock OT1 cells (**Fig. S4C**). These results suggest that SOCS1 decreases the survival of Ag-exp CD8^+^ T cells or prevent the generation of long-lived subsets of CD8^+^ T cells. The former hypothesis is more likely, as tumor-infiltrating sgSOCS1 OT1 cells analyzed 14 days after transfer expressed higher mRNA levels of molecules involved in T cell survival (*Tnfaip3, Bcl2, Il2ra, Il2rb, Jak2)* and cytotoxic/effectors molecules (*Gzmb, Ifngr, Irf1, Fasl, Srgn, Tbx21)* (**Fig. S4D**). Moreover, hallmarks analysis highlighted pathways in tumor-infiltrating sgSOCS1 OT1 cells (*FDR< 0*.*05*), associated with TNFα, IL-2 and IFN-γ responses (**Fig. S4E**). Interestingly, the GSEA of *Socs1*-inactivated OT1 T cells indicates that genes associated with effector functions are more expressed than those implicated in exhaustion (**Fig. S4F**). Targeting *Socs1* in both OT1 and OT2 cells preserved the production of IFN-γ and GzmB in both CD4^+^ and CD8^+^ T cells (**Fig. 4F, G**), while GzmB was increased in CD8^+^ T cells (**Fig. 4F**). Overnight *in vitro* stimulation of sgSOCS1 OT1 cells with titrated SIINFEKL-pulsed DCs led to increased IFN-γ and granzyme B production at high antigen doses after *Socs1* inactivation (**Fig. S4G, H**), showing that *Socs1* actively restrains these cytokines in CD8^+^ T cells. The preserved or increased functionality associated with the increased number of both sgSOCS1 OT2 and OT1 cells led to a much higher number of effector cells at the tumor site (**Fig. 4F, G**), likely explaining the stronger anti-tumor effect of *Socs1* inactivated T cells. Altogether, our results show that SOCS1 has an intrinsic role in the regulation of T cell activation for both CD4^+^ and CD8^+^ T cells.

### Immunotherapeutic potential of SOCS1-edited human CD4^+^ and CD8^+^ CAR T cells

To investigate the therapeutic potential of SOCS1 on human T-cell adoptive transfer, we inactivated SOCS1 gene using Cas9 RNPs in human peripheral blood lymphocytes (PBL) that had been activated and then transduced with a chimeric antigen receptor, encompassing 4-1BB co-stimulatory domains targeting CD19, referred to as 19BBz (**Fig. 5A, B, Fig. S5A, B**). This construct, known to preferentially enhance the survival of CD8^+^ CAR-T cells (CAR8) (Guedan et al., 2018), allowed us to investigate the impact of SOCS1 inactivation on CD4^+^ CAR-T cells (CAR4), which have a limited *in vivo* life-span (Turtle et al., 2016; Yang et al., 2017b). After overnight co-culture with the acute lymphoblastic leukaemia (ALL) FFLuc-BFP NALM6 cell line (NALM6), sgSOCS1 CAR4 and sgSOCS1 CAR8 exhibited a 2-fold higher killing activity (**Fig. S5C**), consistent with the higher levels of effector molecules TNFα, IFN-γ and GzmB that they produced as compared to mock CAR T cells, in three healthy donors (**Fig. S5D, E**). Furthermore, we modelled CAR therapy *in vivo* by injecting 4.10^6^ PBL mock or sgSOCS1-treated (2.10^6^ CAR4 and 2.10^6^ CAR8 cells) in NALM6-infused NOD-scid IL2Rγ^−/−^ (NSG) mice. Seven days after transfer, the number of sgSOCS1 CAR T cells accumulating in bone marrow (BM) was 2-fold higher than that of mock CAR T cells (**Fig. 5C, D**). Reflecting the higher T cell infiltration in the bone-marrow and a more efficient tumor control (**Fig. S5G**), the transcriptomic profiles of sgSOCS1 CAR4 and CAR8 cells at day 7 evidenced upregulation of molecules associated with activation (*FOS, JUND, CD69, SOCS3)*, with long-lived associated factors (*IL7R, PIM1* (Knudson et al., 2017), *TCF7* (Zhou and Xue, 2012) and *KLF2* (Carlson et al., 2006)), resistance to apoptosis (*BCL2L11* (Hildeman et al., 2002) *NDFIP2* (O’Leary et al., 2016)), key regulators of cytotoxic effector functions (*GMZB*, the interferon-induced molecules *GBP5* (Krapp et al., 2016) and *IRF1* and killer associated *NKG7* (Patil et al., 2018)) (**Fig. 5E**).

**Fig. 5.**
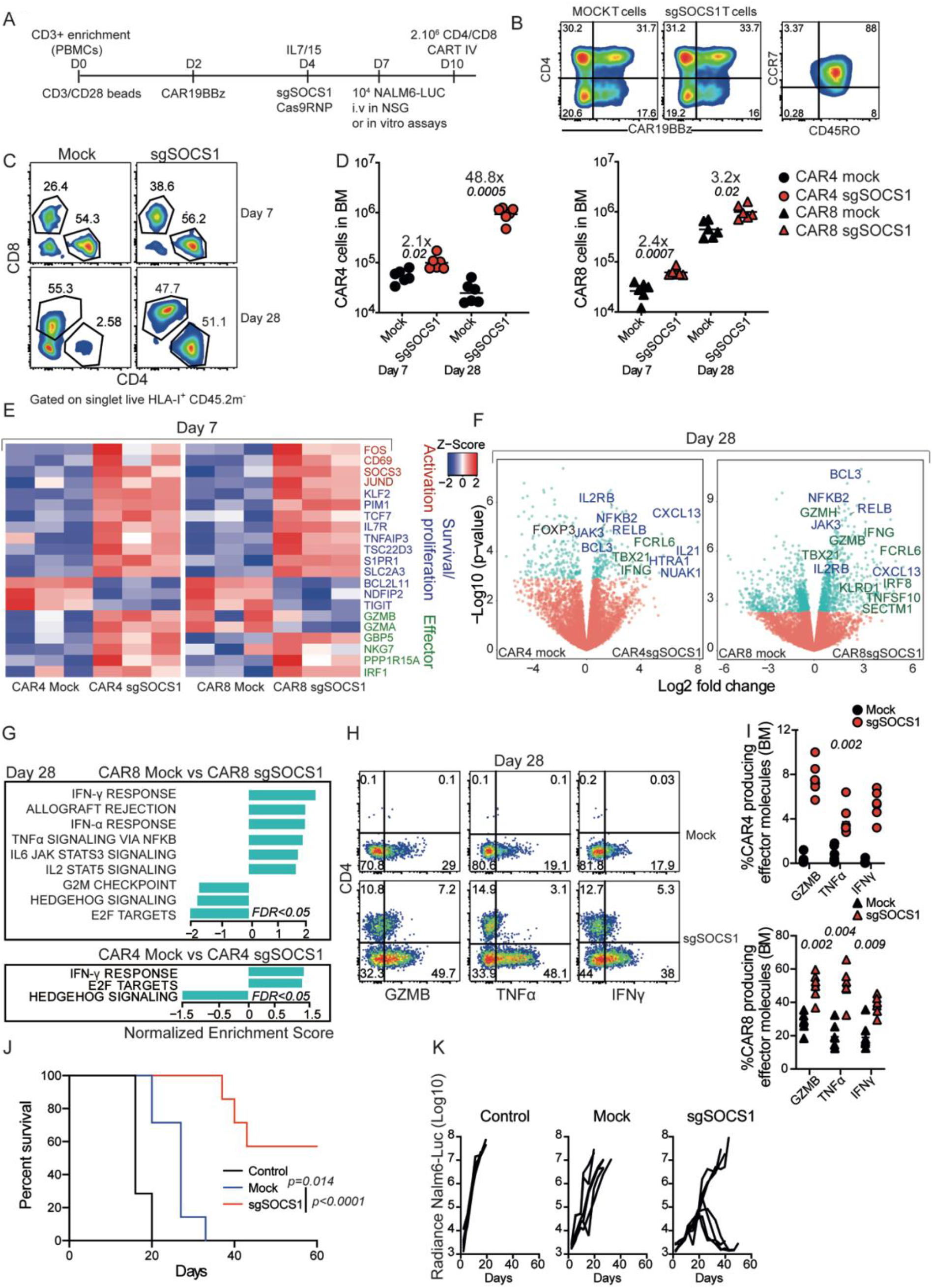
*SOCS1* inactivation restores CAR4 T cell expansion *in vivo* and boosts CAR8 T cell efficacy in controlling B-ALL disease. **(A)** Schematic of CAR-T cell engineering and adoptive T-cell therapy (ATCT) with 2.10^6^ CD4 CAR (CAR4) and 2.10^6^ CD8 CAR (CAR8) T cells of NALM6-Luc-bearing mice. (**B**) CAR expression assessed using CD19/Fc fusion protein and central memory phenotype prior to NSG injection. (**C, D**) Representative flow plots and quantification of bone marrow infiltration with CAR4 and CAR8 mock and sgSOCS1 in NALM6-Luc bearing NSG mice at day 7 and day 28 after transfer, gated on singlets live HLA-I^+^, CD45.2^-^ mouse cells (**E**) Heat map of selected differentially expressed genes (FDR<0.05) between mock and sgSOCS1 CAR T cells related to activation (red), proliferation/survival (blue) and effector functions (green) at day 7 after transfer. (**F**) Differentially expressed genes between mock and sgSOCS1 CAR T cells at day 28 after transfer, with proliferation/survival (blue names) and effector/cytotoxic molecules (green names) highlighted. Transcripts with an FDR value <0.05 are highlighted in light green **(G)** Gene set enrichment analysis of the transcriptional signatures from hallmarks signatures in CAR4/8 sgSOCS1 versus CAR4/8 mock (*n = 6* mice). (**H, I**) Representative flow plots and quantification of effector molecules produced by CAR T cells from infiltrated BM at day 28. (**J, K**) Kaplan–Meier analysis of survival of NSG mice and NALM6-Luc tumor growth after ATCT with 2.10^6^ CAR4/8 mock or 2.10^6^ CAR4/8 sgSOCS1. Data are represented as mean, analyzed by Mann– Whitney U tests or two-way ANOVA, from two independents experiments (*n=3-5* mice/group).

As observed in several studies on CAR-T cell kinetics (Guedan et al., 2018) and CD4/8 CAR-T subset analysis in ALL patients (Turtle et al., 2016; Yang et al., 2017b), CAR8 expanded preferentially over CAR4 in our model. We therefore examined the persistence of sgSOCS1 CAR T-cells, 28 days after transfer. Whereas mock CAR4 declined over time, sgSOCS1 CAR4 and sgSOCS1 CAR8 significantly accumulated in both BM and spleen of NSG mice, correlating with NALM6 rejection (**Fig. S5G, H**). Most strikingly, sgSOCS1 CAR4 expanded to the level of sgSOCS1 CAR8 (**Fig. 5C, D)**. Accordingly, as compared to their mock CAR counterparts in the bone marrow, both sgSOCS1 CAR4 and CAR8 expressed increased levels of cytotoxic/effector-related molecules including *IFNG, FCRL6* (Wilson et al., 2007), *CTSB* (Balaji et al., 2002), *TBX21*, as well as SOCS1-known targets/survival genes such as *IL2RB, JAK3, BCL3* and *CXCL13*, highlighting their tumor reactivity (Li et al., 2019) (**Fig. 5F**). *SOCS1* inactivation led to different transcriptome patterns in CAR4 and CAR8. Transcriptomic analysis of CAR4 evidenced increased expression of pro-survival and self-renewal genes including *IL21*, the insulin growth factor regulator *HTRA1* (Ding and Wu, 2018), and the AMPK-TORC1 metabolic checkpoint *NUAK1* (Monteverde et al., 2018) (**Fig. 5F, G**). This was associated with a proliferation signature represented by E2F targets (**Fig. 5F, G**). CAR8 displayed signs of enhanced cytotoxicity (*GZMB, GZMH, TNFSF10* (TRAIL), Secreted And Transmembrane 1 *SECTM1* (Wang et al., 2012), Killer Cell Lectin Like Receptor D1 *KLRD1* (Li et al., 2019)), some of which were confirmed by flow cytometry analysis (**Fig. 5F, H Fig. S5J**). Contrary to sgSOCS1 CAR4 cells, CAR8 sgSOCS1 expressed lower levels of E2F targets (**Fig. 5G**), suggesting that the higher number of cells found in the BM is related more to survival than proliferation (Ren et al., 2002). While sgSOCS1 CAR cells exhibited a PD1^+^LAG3^+^ phenotype (**Fig. S5J**), implying continuous antigen stimulation, their transcriptional program was more similar to an effector memory than an exhausted signature (Wherry and Kurachi, 2015) (**Fig. S5F**). At late time point (28 days after transfer), not only SOCS1 inactivation led to increased numbers of CAR4 and CAR8 but also to higher cytokine secretion and cytotoxic activity (**Fig. 5H, I Fig. S5I**). Both the increased in numbers and effector functions of the *SOCS1*-inactivated ATCT probably account for their significantly stronger antitumor effects as compared to mock CAR-T cells, with lower tumor load and better survival of NALM6-bearing mice (**Fig. 5J, K**).

## Discussion

Looking for the mechanisms involved in the regulation of CD4^+^ T cell proliferation during an antigenic response, we uncovered SOCS1 as a non-redundant signaling node, leading to a negative feedback loop downstream of TCR and lymphokine signaling. SOCS1 appears to actively restrain T cell proliferation, survival and effector functions during an antigenic immune response. Inactivating *Socs1* evidenced different effects on CD4^+^ and CD8^+^ T cells: it greatly improved CD4^+^ T cell proliferation and survival while it mostly increased the effector function of CD8^+^ T cells with a small effect on cell survival. *Socs1* inactivation in T cells specific for tumor antigens resulted in enhanced anti-tumor activity after adoptive transfer in both mouse and human models. This may have important therapeutic implications.

In two different TCR-Tg models (Marilyn and OT2), we demonstrated that SOCS1 is a major intrinsic negative regulator of Ag-exp CD4^+^ T-cell expansion *in vivo*. We report the same findings in these two CD4^+^ T-cell models exhibiting distinct avidities, mode of secretion (Robinson et al., 1986) and using different type of antigenic stimulations such as DC-peptide stimulation or tumor challenge. Altogether, this highlights the generalizable aspect of our discovery, operating for all CD4^+^ T cells. Surprisingly, our *in vivo* genome-wide positive screen evidenced only one hit, *Socs1*. It is highly probable that the genes necessary for *in vitro* growth and survival were missed as we used a constitutive CRISPR/Cas9 system. In addition, the restored proliferation of antigen experienced Marilyn cells by blocking both IL2 and IFN-γ pathways in our model, suggest a genetic redundancy and compensation between inactivated receptors that could not be revealed by our screening strategy.

Our data suggest that cytokine sensing play a role in impairing CD4^+^ T cells immunity after Ag re-exposure/chronic stimulation. This paradoxical cytokine-mediated suppression of CD4^+^ T cells has already been described, when blocking chronic IFN-I signaling during persistent infection enhanced CD4^+^ T cell-dependent virus clearance (Teijaro et al., 2013; Wilson et al., 2013). SOCS1 may be responsible for the so-called activation induced cell death (AICD), where IL-2 (Lenardo, 1991) or IFN-γ (Berner et al., 2007) provided too early after antigen stimulation leads to apoptosis of CD4^+^ T cells (Majri et al., 2018). Hence, we observed that SOCS1 prevented the expression of genes involved in resistance to apoptosis, such as *Bcl2, Bcl3, Tnfaip3, Hopx* (Albrecht et al., 2010). SOCS1 also appears to selectively regulate the proliferation of CD4^+^ T cells as compared to CD8^+^ T cells *in vivo* by inhibiting, both in human and murine CD4^+^ T cells the expression of E2F targets, which are key regulators of cell cycle progression (Zhu et al., 2001). Thus, targeting SOCS1 improves CD4^+^ T cells survival and proliferation by rendering them insensitive to out of sequence lymphokine-induced cell death. This phenomenon has been described for SOCS3, another member of SOCS family, which is involved in the impairment of human and murine CD4^+^ T cells *in vivo*, after cytokine pre-exposure (Sckisel et al., 2015). However, SOCS3 expression is associated with Th2 lineage commitment, while SOCS1 is involved in Th1 differentiation (Egwuagu et al., 2002).

As SOCS1 negatively regulates Ag-exp CD4^+^ T-cell capacity to produce several cytokines essential for anti-tumor immunity (Dobrzanski, 2013), we explored the direct and indirect impact of *Socs1* deletion on adoptively transferred antitumor CD4^+^ T cells. Targeting SOCS1 also increases Ag-exp CD4^+^ T-cell polyfunctionality *in vivo*, enhancing their lymphokines secretion, in particular IFN-γ in the TdLN (Fig. 3) and at the tumor site (**Fig. 4, Fig. 5**). This may account for the higher number of active CD8^+^ T cells and NK cells infiltrating the tumor (**Fig. 3I, J; Fig. 4F,G**). The increased infiltration and persistence of functional CD4^+^ T cells could participate to tumor rejection by helping CD8^+^ T cell priming or migration to the tumor bed, recruiting innate cells or directly killing tumor cells (Borst et al., 2018). Thus, both murine and human CD4^+^ T cells targeted for SOCS1 exhibit an increased expression of the cytotoxic molecule GZMB at the tumor site (**Fig. 3, Fig. 5**). The acquisition of such cytotoxic features by CD4^+^ T cells have been recently associated with a Blimp-1-dependent IL-2 autocrine stimulation (Śledzińska et al., 2020). Hence, through both extrinsic and intrinsic mechanisms (Zander et al., 2019), the transfer of *Socs1*-deleted tumor specific CD4^+^ T-cell improve the magnitude of the endogenous antitumor immune response as well as infiltration by cytotoxic CD4^+^ T cells, which act in concert toward tumor eradication.

With no effect on CD8^+^ T cell division and little impact on their survival *in vivo*, SOCS1 significantly impedes CD8^+^ T cell cytotoxicity. This could be the result of enhanced sgSOCS1 CD4+ helper function as both CD4^+^ and CD8^+^ T cells are co-transferred. However, Marilyn sgSOCS1 do not increase the cytotoxic capacity per cell basis of CD8^+^ and NK cells from the endogenous compartment. Furthermore, we demonstrated *in vitro* that SOCS1 actively restrained the TCR-induced capacity to produce IFN-γ and GZMB of CD8^+^ T cells. Altogether, this demonstrated that SOCS1 inactivation per se increased the cytotoxic potential of T cells. This effect could be mediated by IRFs, which are central amplifier circuits downstream IFN-I and IFN-II signaling (Michalska et al., 2018), regulating the expression of both IFN-γ and granzyme B (Guinn et al., 2016). In addition to IRF4, it appears that other IRFs family members, including IRF1 and IRF8 (**Fig. S4, Fig. 5**) are modulated by SOCS1.

With an improved persistence *in vivo*, SOCS1 targeted CD4^+^ T cells are probably subjected to chronic stimulation that might lead to anergy and Tregs conversion (Alonso et al., 2018). However, SOCS1 is essential for the maintenance of Foxp3 expression and for Tregs suppressive functions *in vivo* (Takahashi et al., 2011, 2017). Accordingly, *Socs1-*inactivated Marylin CD4^+^ T cells display enrichment of conventional T cells markers as opposed to Tregs genes (**Fig. S3C**). Moreover, we noticed a decreased gene expression of *FOXP3* in sgSOCS1 CAR4 as compared to mock CAR4 at late time point (**Fig. 5G**). Altogether, it seems that targeting SOCS1 in CD4^+^ T cells may prevent them to convert into Tregs.

The forced expression of cytokine-encoding genes or construct containing a JAK/STAT signaling domain in CD8^+^ CAR-T cells improves their persistence and antitumor effects *in vivo*, highlighting the importance of signal 3 for CAR-T cell functions (Kagoya et al., 2018; Markley and Sadelain, 2010; Quintarelli et al., 2007). Here, we demonstrate that inactivating a major inhibitor of cytokines signaling in CAR-T cells also enhance their therapeutic potential and most importantly selectively affect CD4^+^ and CD8^+^ CAR-T cells. This has major relevance to design and potentiate the next generation of adoptive T cells therapies for cancer and viral infections with an optimized composition and improved efficacy. However, the reduced control in cytokine signaling could potentially increase the risk for cytokine release syndrome (CRS), especially since SOCS1 has been shown to regulate IL6 signaling (Diehl et al., 2000). This issue might nevertheless be addressed using an inducible and reversible gene inactivation for clinical translation (Lucibello et al., 2020).

Our findings demonstrate the feasibility of interrogating *in vivo* genome-wide mutated primary CD4^+^ T cells, applicable to further study CD4^+^ T-cell or regulatory T-cell plasticity and functions. This work underlies the relevance of understanding CD4^+^ T cell biology for the development of effective T cell-mediated therapies. We unravel the importance of signal 3 regulation in CD4^+^ T cell biological functions and identified a major intracellular checkpoint critical for the magnitude, duration and quality of T cell immune responses, that may prove efficacy in clinics.

## Acknowledgments

We thank Lorenzo Galluzzi for his general comments, Nina Burgdoff and Sheila Lopez for the technical support they provided with *in vivo* and CAR-T cells experiments. We thank the zootechnicians of the mouse facility plateforme of the Institut Curie as well as the flow cytometry core. We thank Audrey Rapinat from the Genomics Platform of the translational research department at Institut Curie. We thank Jessie Thalmensi for providing advices concerning the MB49 tumor model and Christel Goudot for coordinating the NGS sequencing analysis. High throughput sequencing has been performed by the ICGex NGS platform of the Institut Curie (Sonia Lameiras, Virginie Raynal, Patricia Legoix) supported by the grants ANR10EQPX03 (Equipex) and ANR10INBS0908 (France Génomique Consortium) from the Agence Nationale de la Recherche (“Investissements d’Avenir” program), by the Canceropole Ile de France and by the SiRICCurie program. Laurie Menger was supported by a Marie-Sklodowska Curie individual fellowship (grant 743435) from the European Commission, INSERM and Agence Nationale de la Recherche (ANR jeune chercheur JCJC 2018). Olivier Lantz was supported by the Institut National de la Santé et de la Recherche Médicale, Institut Curie, ARC fundation, fondation Trouver et Chercher.

## Author Contributions

Conceptualization: L.M; O.L; Methodology and Investigation: A.SDG, S.M, F.L, J.F, A.D, M.S, M.D, J.T, P.G; Writing – Original Draft, L.M; Writing – Review & Editing: J.H, DC.R, M.S, G.S, S.A, O.L, L.M.

## Methods

### Lead Contact

Further information and requests for resources and reagents should be directed to and will be fulfilled by the Lead Contact, Laurie Menger

### Materials Availability

This study did not generate new unique reagents.

### Data and Code Availability

The affymetrix and RNAseq data supporting this study have been deposited in the GEO database under the accession number GSE154794.The SRA database accession number for the screens analysis is PRJNA639469 (Temporary Submission ID: SUB7577588) and will be accessible at https://www.ncbi.nlm.nih.gov/sra/PRJNA639469.

### Experimental model and subject details

#### Cell lines and mice

B16-OVA and MB49 cell lines, kindly provided by E. Piaggio and C. Théry, FFLuc-BFP NALM6 (NALM6) cell line, provided by O. Bernard were maintained in RPMI-1640 supplemented with 10% FBS. CD45.1 and CD45.2 female Marilyn TCR-transgenic Rag2^−/−^ mice, specific for the HY male antigen were crossed to Rosa26-Cas9-EGFP knock-in mice (026179, Jackson lab). Thy1.1 and Thy1.2 OT-II TCR-transgenic Rag2^−/−^ mice, CD45.1 female OT-I TCR-transgenic Rag2^−/−^ mice, specific for OVA and female, male NOD-scid IL2Rγ ^−/−^(NSG) mice were also used in this study. Female C57BL/6 mice were purchased from Charles River Laboratories (L’Arbresle, France). All experiments were conducted with 6-12 weeks old mice, in an accredited animal facility by the French Veterinarian Department following ethical guidelines, approved by the relevant ethical committee (*AP AF1S#6030-20 16070817147969 v2*, authorisation #XX DAP 2017-023).

#### Cell culture and adoptive transfers

Naive CD4^+^ T cells were obtained from peripheral lymph nodes of Marilyn or OT-II mice. Antigen experienced CD4^+^ T cells were generated *in vitro* by priming lymph nodes and splenocytes of CD45.1 Marilyn mice or Thy1.1 OT-II mice with respectively 10nM Dby (NAGFN-SNRANSSRSS, Genscript) and 5μM OVA_II_ peptide (InvivoGen). IL-2 (10ng/mL), IL-7 (2ng/mL) (Peprotech) were added starting at day 4 and every 3 days in complete RPMI-1640 supplemented with 10% FBS and 0.55 mM β-mercaptoethanol, while resting for 6-10 days. Ag-exp OT-I cells from lymph nodes and spleen were cultured with 0.5μM SIINFEKL (InvivoGen) and maintained with IL15 (50ng/mL) (Peprotech) every two days. T cells were labeled with 5μM CFSE (Invitrogen) in PBS for 8 minutes at 37°C. For *in vivo* GS screen, 4.10^6^ naïve CD45.2 Marilyn CD4^+^ T cells were transferred and footpad vaccinated with 4.10^6^ Dby-loaded-LPS-matured bone marrow derived-dendritic cells (BMDCs). Seven days later, 12.10^6^ library-transduced or 12.10^6^ Mock-transduced CD45.1 Cas9-Marilyn cells were injected intravenously and mice were at the same time footpad-vaccinated with 4.10^6^ Dby-loaded-LPS-matured BMDCs. For validation experiments, a first cohort of 10^6^ naïve CD45.2 Marilyn or Thy1.2 OT-II cells was transferred into CD45.2 B6 hosts footpad vaccinated with 10^6^ peptide-loaded-LPS-matured BMDCs. After 7 days, a second cohort of either 10^6^ naïve CD45.1 Marilyn, Thy1.1 OT-II cells or 2.10^6^ Ag-exp CD45.1 Marilyn, 2.10^6^ Thy1.1 OT-II CD4^+^ T cells were injected and mice were footpad-vaccinated with 10^6^ peptide-loaded-LPS-matured BMDCs. The number of injected cells as a second cohort is based on CD62L positivity, reflecting the capacity of naïve and memory (Ag-exp) CD4 T cells to similarly home to the LN. BMDCs were generated by 10 days culture in complete IMDM containing 20ng/ml of GM-CSF (Peprotech) and maturation was induced by a 20-hour treatment with 1ug/mL lipopolysaccharide (Sigma-Aldrich), pulsed with 50nM Dby or 20μM OVA_II_ peptide for 2hours. Mice were treated with blocking antibodies from Bioxcell, including isotypes control rat IgG2b (clone LTF2), IgG2a (clone 2A3), anti-mouse CD122 antibody (clone TM-Beta1), anti-mouse IFN-γR (clone GR-20), intraperitoneally on day 7, 11 and day 11 after ACT (10 mg/kg).

For adoptive cell therapies, female C57BL6 host were subcutaneously implanted with either 1.5.10^6^male bladder MB49 tumor cells or 4.10^5^ B16-OVA melanoma cells. At day 10 for the MB49 model and on day 7 for B16-OVA, 10^6^ Marilyn CD4^+^ T cells or 2.10^6^ OT-I and 2.10^6^ OT-II cells were adoptively transferred into tumor-bearing mice (n = 4-6/group). Mice were sacrificed when the tumors exceeded 15 mm in diameter for the B16-OVA model.

Peripheral blood mononuclear cells (PBMCs) from healthy donors were isolated by density gradient centrifugation. T lymphocytes were purified using the Pan T cell isolation kit (Miltenyi Biotech) and activated with Dynabeads Human T-Activator CD3/CD28 (1:1 beads:cell) (ThermoFisher) in X-vivo 15 medium (Lonza) supplemented with 5% human serum (Sigma) and 0.5 mM β-mercaptoethanol at density of 10^6^ cells/mL. 48 hours after activation, T cells were transduced with lentiviral supernatants of an anti-CD19(FMC63)-CD8tm-4IBB-CD3ζ CAR construct (rLV.EF1.19BBz, Flash Therapeutics) at MOI 10. Two days later, the CD3/CD28 beads were magnetically removed, CAR T cells were electroporated with Cas9-ribonucleoproteins (Cas9-RNP) and maintained in X-vivo supplemented with IL7 (5ng/mL) and IL15 (5ng/mL). Six days after electroporation, CD4^+^ and CD8^+^ CAR-T cell were separated using CD8^+^ T Cell Isolation kit (Miltenyi) for mutagenesis quantification on gDNA and western blot analysis of SOCS1 expression.

Male or female 8–12-week-old NSG mice were injected with 2.10^5^ NALM6 cells intravenously by tail vein injection. Three days later, 2.10^6^ CAR T cells were administered intravenously by tail vein injection (day 0). Tumor burden was measured by bioluminescence imaging using the Lumina IVIS Imaging System (PerkinElmer). Mice were sacrificed when the radiance was > 5.10^6^ [p/s/cm≤/sr].

#### Cytotoxicity assays

The cytotoxicity of T cells transduced with a CAR was determined by co-culturing in triplicates at the indicated *E*/*T* ratio, CAR T cells (Effectors) with Nalm6 cells (Targets) in a total volume of 100 μl per well in X-vivo medium. The maximal luciferase expression (relative light units; RLUmax) was determined with target cells alone plated at the same cell density. 18h later, 100μl luciferase substrate (Perkin Elmer) was directly added to each well. Luminescence was detected using a SpectraMax ID3 plate reader (VWR). Lysis was determined as (1 − (RLUsample)/(RLUmax))× 100.

#### Antibodies and Flow cytometry analysis

Lymph nodes cells, splenocytes and tumor samples enriched on a density gradient medium (Histopaque, Sigma) were incubated with murine antibodies (Key Resources Table). Human cultured cells, bone marrow cells and splenocytes from NSG mice cells were stained with the indicated Abs or soluble protein: fluorochrome-conjugated antibodies specific for human (Key Resources Table). The intracellular staining was performed either with intracellular staining permeabilization wash buffer (BD Bioscience) or Foxp3 kit (eBioscience). CAR expression was assessed using 9269-CD-050 Recombinant Human CD19 Fc Chimera Protein (Bio Techne), at 4°C for one hour, at 1/100 dilution. Viability was evaluated using Fixable Viability Dye eFluor 780 (eBioscience) or Aqua Live dead (Thermo Fisher). Re-stimulation was performed with 20ng/mL of PMA (Sigma), 1μM of ionomycin (Sigma) and BD Golgi plug for 4 hours at 37°C. Cell Sorting Set-up Beads (Life Technologies) were used to quantify and normalized cell number between samples and experiments. Stainings were performed in a blocking solution: 5% FCS, and 2% anti-FcR 2.4G2, and samples acquired on a LSRII/ Fortessa (BD) and analyzed with FlowJo software (V10, Tree Star). Cell sorting was performed on ARIAII (BD).

#### Western blot analysis

T cells (2.10^6^) were lysed using RIPA lysis buffer (Thermofisher) and 1X Protease Inhibitor Cocktail (Sigma). Cell debris were removed by centrifugation at 14,000 rpm for 15 min at 4°C and 20-40μg of proteins from the supernatant were separated using SDS-PAGE and transferred to a PVDF membrane. SOCS1 and β-actin (loading control) were visualized using monoclonal antibodies anti-SOCS1 (1μg/mL) (ab62584; Abcam), anti-Actin mouse (Millipore, clone C4), HRP-anti-Rabbit IgG1 (Cell Signaling Technology). HRP-anti mouse IgG (Cell signaling) on Chemidoc Touch Imaging system (Biorad). Signal instensity was quantified with ImageJ software.

#### Genome-wide CRISPR-Cas9 screens

The lentiviral gRNA plasmid library for genome-wide CRISPR-Cas9 screen (Mouse Improved Genome-wide Knockout CRISPR Library v2, Pooled Library #67988#) and mock vector (#67974) was obtained from Addgene. The library was amplified following the protocol provided by Addgene. Briefly, 4×25ul of NEB 10-beta Electrocompetent *E. coli* (NEB, cat. no. C3020K) were electroporated with of 4×10 ng/µl and cultured in 4×500mL of ampicillin-treated Luria-Bertani (LB) incubate at 37 °C overnight with shaking. The plasmids were extracted with 12 columns of EndoFree plasmid Maxi kit (Qiagen). To prepare the virus library, 293T cells at low passage (<7) in 20cm dish (X15) were transfected with 11 μg of gRNA library, 11 μg of psPAX2 and 2.5 μg of pVSV-G. Twenty-four hours after transfection, the medium was changed to DMEM-1% BSA, collected at 48h, 60h and 72h, then centrifuged, filtered through 0.45uM PVDF membranes (Millipore), concentrated using Amicon Ultra 15ml centrifugal filters (Merck) and used fresh. One day before T cells transduction, CD4^+^ T cells are enriched using MagniSort Mouse CD4^+^ T cell Enrichment Kit (Thermofisher scientific) and seeded at a density of 1,5.10^6^ cells/ml with ½ fresh medium and ½ culture medium supplemented with IL-2 (10ng/ml), IL-7 (2ng/ml). Cells are spinfected for 90min, at 32°C, 900g with 10ug/ml of protamine sulfate (Sigma) and 8ug/ml of DEAE-dextran (Sigma). The volume of the lentivirus library used is the one required for achieving an optimal transduction efficiency, MOI of 0.3 after 5 days selection with 5ug/ml of puromycin (Sigma). CFSE^hi^ and CFSE^lo^ Cas9-CD45.1 Marilyn CD4^+^ T cells were sorted and their gDNA extracted using 10μl of lysis buffer-AL (Qiagen-DNeasy blood and tissue kit), 1μl proteinase K (Qiagen), followed by 30 min incubation at 56°C, 30 min incubation at 95°C and resuspension in 20μl of ddH20 on ice. The gRNAs were amplified by a two-step PCR method using the Herculase II Fusion DNA Polymerase (Agilent). For the first step PCR, all the gDNA extracted is used to perform approximately 30×50-μl PCR reactions with the forward primer 50bp-F and the reverse primer 50bp-R (Key Resources Table); the PCR program used is 94 ° C for 180 s, 16 cycles of 94 ° C for 30 s, 60 ° C for 10 s and 72 ° C for 25 s, and a final 2-min extension at 68 ° C. Products of the first-step PCR are pooled, purified with Ampure XP (Agencourt) and quantified using the dsDNA HS assay kit. Three 50-μl PCR reactions were performed with the forward primer Index-F and one of the reverse primers (Index-R1 to R6). The PCR program used is 94 ° C for 180 s, 18 cycles of 94 ° C for 30 s, 54 ° C for 10 s and 72 ° C for 18 s, and a final 2-min extension at 68 ° C. Products of the second-step PCR reactions were purified and analysed with Caliper Labchip for DNA samples (HT DNA High Sensitivity LabChip Kit; Perkin Elmer) prior to sequencing with the Miseq or HiSeq2500 instrument for the library representation (Illumina). The DNA quality was assessed and quantified using an Agilent DNA 1000 series II assay and a Qubit fluorometer (Invitrogen). Sequencing was performed with a 10% Phix control, using the 25-bp single-end sequencing protocol preceded by 23 dark cycles to mark the repetitive structure of the target region.

#### Bulk mRNA Sequencing and Analysis

Between 10^4^ and 3.10^4^ murine and human T cells were sorted from lymph nodes and tumors in TCL buffer (Qiagen) with 1% of β-mercaptoethanol. Total RNA was purified using the Single Cell RNA purification kit (Norgen) according to the manufacturer’s instructions, including a step of DNAse treatment (Qiagen). The RNA integrity number was then evaluated with an Agilent RNA 6000 pico kit. cDNA synthesis and Illumina-compatible libraries were generated from total RNA (0,25-10ng) by Next Generation Sequencing platform of the Institut Curie, using SMARTer Stranded Total RNA-Seq Kit-Pico Input Mammalian according to manufacturer’s instructions. Libraries were then sequenced on an Illumina NovaSeq-S1 using 100bp paired-end mode (OR HiSeq - Rapid Run - PE100). FASTQ files were mapped to the reference genome hg19 (human) or mm10 (mice) using Hisat2 and counted by featureCounts from the Subread R package to produce read count tables. EdgeR was then used to normalize read counts and gene with expression > 0.5 cpm in at least three replicates were kept for subsequent analysis. Differential gene expression was performed with limma-voom R package. The fgsea R package was used to compute the enrichment scores. For Affymetrix analysis, gene expression was conducted using Mouse Clariom D chip (Thermo Fisher). RNA samples were amplified with Ovation Pico WTA System v2 (Nugen) and labeled with Encore biotin module (Nugen). Array were hybridized with 5 µg of labeled DNA and assayed on a GeneChip Scanner 3000 7G (Affymetrix). Raw data were generated and controlled with Expression console (Affymetrix) at the Institut Curie Genomic facility.

#### Genome-wide data processing

FASTQ files obtained after sequencing were demultiplexed using the HiSeq Analysis software (Illumina). MAGeCK (Li et al., 2014) count command was then used to generate per-sgRNA read count table by matching single-end reads with sgRNA sequences from the genome-scale sgRNA Yusa library (Koike-Yusa et al., 2014). Before mapping, the library was first cleansed of (i) all sgRNA that did not map the reference genome (here mm10) and (ii) all sgRNA that mapped multiple spot in the reference genome (multihits). Redundant sgRNA were merged. A normalizing factor for each sample was then calculated using Trimmed Mean of M-values (TMM) method implemented in edgeR R package (Robinson and Oshlack, 2010) Normalized counts were filtered for low expressed sgRNA (keeping only sgRNA with at least 4 count per million in 3 samples) and transformed to log2-counts per million using voom implemented in limma R package. Differential expression of each sgRNA was calculated using lmFit function in limma using the high and low CFSE cell fraction from each screen. For each sgRNA, enriched and depleted p-values were computed using one-tailed paired Student’s t-tests. From these, Robust Rang Aggregation (RRA) score (10.1093/bioinformatics/btr709) for each gene was computed among multiple sgRNAs (n=5) of each gene and gene-level related p values and corresponding adjusted p-values [False Discovery Rates (FDR)] were obtained using a permutation test with 1,000,000 iterations with same size randomized gene sets. Finally, graphical representation of genes according to their enriched p value and median log fold change of sgRNA supporting the RRA score was done.

#### Cas9-RNP validations

1 µl Oligos crRNA (100nM) and 1µl tracrRNA (100nM) (Key Resources Table) for murine T cells and 1 µl Oligos crRNA1 + 1 µl Oligos crRNA2 +1 µl Oligos tracrRNA for human T cells were annealed at 95°C for 5min and incubated at room temperature 10 min with 10μg S.p Hifi Cas9 Nuclease V3. 2.10^6^ T cells were resuspended in 20 µl of nucleofection solution with 3 µl or 4 µl RNP and transferred to Nucleofection cuvette strips (4D-Nucleofector X kit S; Lonza). Murine T cells were electroporated using the DN110 program of 4D nucleofector (4D-Nucleofector Core Unit: Lonza, AAF-1002B), human CAR T cells using the program E0115. T cells were then incubated at 32°C for 24 to 48 hours to increase the mutagenesis efficacy (Doyon et al., 2010), prior to resuspension in supplemented fresh medium. Murine CD4^+^ T cells were maintained in complete RPMI with IL2 (10ng/mL) and IL-7 (2ng/mL). Human T cells were maintained in X-Vivo with 5% human serum and IL7 (5ng/mL) and IL15 (5ng/mL). Locus-specific PCRs (Key Resources Table) were performed on genomic DNA and frequencies of NHEJ mutations were assessed by sequencing (Eurofins, Mix2seq) and TIDE analysis (https://tide.deskgen.com).

#### Statistical Analysis

One-way ANOVA, two-way ANOVA, or Mann–Whitney non-parametric test with *p*< 0.05 were performed using Prism 8.0 software (GraphPad). Multiple comparisons were corrected with the Bonferroni coefficient and Kaplan–Meier survival curves were compared with the log-rank test.

## Supplemental Information titles and legends

**Fig. S1. Establishing and validating *in vivo* Genome-wide pooled CRISPR screens in CD4 T cells, related to figure 1**. (**A)** Percentage (gated on singlets live CD45.1^+^ CD4 T cells) of naïve and Antigen-experienced (Ag-exp) Marilyn-Cas9 cells proliferation *in vivo* 14 days after footpad vaccinations. (**B**) CD45.1 Marilyn (Vβ6^+^) CD4 T cell Cas9 expression phenotype (red) after crossing CD45.1 Marilyn TCR (blue) transgenic Rag2^−/−^ mice to Rosa-26-Cas9-EGFP knock-in mice and activation (CD44), homing (CD62L) markers in naïve and in Ag-exp Marilyn CD4 T cell (*in vitro* priming and resting) prior to injection. (**C**) Representative plots showing transduced Ag-exp Marilyn-Cas9 cell viability and BFP reporter expression before and after puromycin selection (5μg/mL for 4 days). (**D, E**) Deep-sequencing analysis (Hiseq 300 million reads) of the gRNAs in the lentiviral plasmid DNA library (D) and in the genomic DNA of 45.10^6^ transduced Marilyn-Cas9 cells (E). The percentage of sgRNA with read count < 10 are mentioned in red. (**F**) SORTing strategy of CFSE^lo^ and CFSE^hi^ Marilyn-Cas9 library-transduced after *in vivo* selection. (**G**) Box-dot plot of overall sgRNA library representation in CFSE^hi^ and CFSE^lo^ sorted population ex-vivo (*n=3-6* mice/group). **(H)** Dot plot confirming sgSOCS1 enrichment (*n* = 5 sgRNAs per gene) in CFSE^lo^ subset of CD45.1-library-transduced CD4 T cells compared to the CFSE^hi^ subset without the first cohort (weak selection), from three independent GW screens, two-tailed paired Student’s t-test. (**I**) Tide analysis showing the percentage of NHEJ-mutations in the genomic DNA (gDNA) of Ag-exp Marilyn and OT2 cells, 4 days after electroporation with sgSOCS1. **(J)** SOCS1 protein expression in Ag-exp Marilyn and OT2 cells 6 days after electroporation by western blot analysis. (**K**) Representative plots and percentage (gated on singlets live CD90.1^+^ CD4 T cells) of proliferating Ag-exp Mock or sgSOCS1 OT2 cells during a recall response, at day 14. Mice were injected with 2.10^6^ CD4 T cells IV and primed with 10^6^ peptide–pulsed LPS-matured DCs at day 0 and day 7. (**L**) Representative flow plots showing the percentage of Marilyn and OT2 Tregs (CD25^+^FOXP3^+^) in C57BL/6 mice transferred and vaccinated from Fig.1I and Fig.S1K, at day 14. Data are shown as mean, analyzed by Mann–Whitney U tests from two (K) or three independent experiments (H), *n=2-6* mice/group.

**Fig. S2. Validating SOCS1 as negative feedback node integrating/regulating several lymphokines signals, related to figure2**.

**(A)** Representative flow and percentage of CD25 expression in naive and Ag-exp Marilyn cells in the course of an ongoing immune response. **(B)** Representative flow plots showing naive and Ag-exp Marilyn cells producing IL2 and IFN-γ in an ongoing immune response. **(C)** Representative flow plots (percentage highlighted are from singlets live CD45.1^+^ CD4 T cells) and quantification of 10^6^ Ag-exp Marilyn IFNγ-R^−/−^ cells expansion *in vivo* during a CD4^+^ recall response, in the presence of blocking antibodies (200μg) injected intraperitoneally at day 7, day 9, day 11: isotypes, anti-IL2Rβ. (**D**) Flow cytometric evaluation of CD69, CD25, IRF4 and expressions in sgSOCS1 Ag-exp OT2 cells compared to Mock cells after overnight co-culture with peptide–pulsed LPS-matured DCs *in vitro*, in the absence of cytokine. (**E**) Representative flow plots showing CD25 and IRF4 expressions in Marilyn and OT2 cells mock or sgSOCS1 after overnight coculture with DCs loaded with increasing doses of Dby. (**F**) Flow plots and percentage of IFN-γ-, TNFα- and IL-2-producing Mock or sgSOCS1 OT2 cells. Data are shown as mean or mean ± SD, analyzed by Mann–Whitney U tests or two-way ANOVA (D), from two independent experiments *n=2-3* mice/group.

**Fig. S3. *Socs1* gene inactivation improves Marilyn adoptive cell therapy of male bladder tumors MB49**.

(**A**) Differentially expressed genes in Tumor draining lymph node (TdLN)-infiltrating CD45.1 Marilyn sgSOCS1 cells compared to Marilyn mock cells. Transcripts with an FDR value <0.05 are highlighted in light green. (**B**) Representative flow plots and quantification of CD45.1^+^ Marilyn cells in the blood of MB49-bearing C57BL/6 mice at day 25. (**C**) Absolute number of polycytokine producing Marilyn cells in the TdLN at day 7 after transfer. (**D**) Genes uniquely downregulated in naïve vs effector-memory CD4 T cells (left panel, GSE11057) and upregulated in Tregs vs Tconv in lymph node (LN, right panel, GSE37532) were evaluated in Marilyn sgSOCS1 versus Marilyn mock in TdLN (*n = 6* mice) using Gene set enrichment analysis. NES: normalized enrichment score. **(E)** Representative flow plots and quantification of functionally active (PD1^+^, Tim3^+^) CD8 and NK cells from the endogenous compartment, infiltrating the tumor at day 7. Data are shown as mean, analyzed by Mann–Whitney U tests from two independent experiments, *n=4-6* mice*/*group.

**Fig.S4. *Socs1* gene inactivation in Ag-exp OT1 and OT2 cells, related to figure 4**.

**(A)** TIDE analysis showing the percentage of NHEJ-mutations in the gDNA of OT1 cells, 4 days after electroporation with sgSOCS1. **(B)** SOCS1 protein expression in OT1 cells 6 days after electroporation by western blot analysis. (**C**) Representative plots, quantification and phenotype of OT2 and OT1 cells in 50μL of blood from B16-OVA challenged mice at day 60 after tumor challenge. (**D**) Differentially expressed genes in tumor-infiltrating CD45.1 OT1 sgSOCS1 cells compared to OT1 mock cells, 14 days after transfer. Transcripts with an FDR value <0.05 are highlighted in light green. (**E**) Gene set enrichment analysis (GSEA) of selected Hallmarks transcriptional signatures (MSigDB) with an FDR value < 0.05 in Ag-exp OT1 sgSOCS1 versus Ag-exp mock OT1 cells in the tumor (*n = 3* replicates from 2 pooled mice). (**F**) Genes uniquely upregulated in effector CD8 T cells vs exhausted CD8 T cells (GSE9650) were assayed in OT1 sgSOCS1 cells vs OT1 Mock cells in tumor at day 14. (**G, H**) Effector molecules produced by OT1 sgSOCS1 and OT1 mock cells after overnight co-culture with DCs-loaded with increasing doses of SIINFKEL peptide *in vitro*, from two independent experiments, analyzed by two-way ANOVA. Data are shown as mean ± SD analyzed by Mann–Whitney U tests from two independent experiments, *n=4-6* mice/group.

**Fig. S5. *SOCS1* inactivation improves CAR4 and CAR8 expansion, persistence and functional activity, related to figure 5**.

**(A)** TIDE analysis showing the percentage of NHEJ-mutations in the gDNA of human CD4 (CAR4) and CD8 CAR T cells (CAR8), 4 days after electroporation with sgSOCS1. **(B)** SOCS1 protein expression in CAR4 and CAR8 cells 6 days after electroporation by western blot analysis. (**C**) Cytotoxic activity using an 18 h bioluminescence assay, using firefly luciferase (FFL)-expressing NALM-6 (NALM6-Luc) as targets cells (*n =3* healthy donors). (**D, E**) Flow plots and quantification depicting CAR4 and CAR8 mock and sgSOCS1 effector molecules produced after overnight co-culture with NALM6-Luc ALL cells (n=3 donors). (**F**) Genes uniquely downregulated in naïve vs effector CD8 T cells (top panel, KAECH) and upregulated in effector CD8 T cells (Teff) vs exhausted CD8 T cells (Texh) (bottom panel, GSE41867) were assayed in CAR8 sgSOCS1 versus CAR8 mock (*n = 6* mice) using Gene set enrichment analysis. (**G**) Representative flow plots (day 28) and quantification of NALM6-Luc cells infiltrating NSG mice 7 days and 28 days after CAR transfer, gated on singlets live HLA-I^+^, CD45.2^-^ mouse cells (2.10^6^ mock CAR4/CAR8 or 2.10^6^ sgSOCS1 CAR4/CAR8). (**H**) Mock CAR4/CAR8 and sgSOCS1 CAR4/CAR8 infiltration in NSG mice spleen, 28 days after transfer, gated on live HLA-I^+^, CD45.2^-^ mouse cells. (**I**) Number of CAR4 and CAR8 producing effectors molecules from the bone marrow at day 28 *ex vivo*. (**J**) Representative flow plots and quantification of negative checkpoints expressed by CAR4/CAR8 cells in the bone marrow of NALM6-Luc transferred NSG mice at day 28 (*n = 6* mice). Data are represented as means or mean ±SD, analyzed by Mann–Whitney U tests, from three independent experiments (A-E) or two independents experiments (F-J).

